# A novel brain penetrant PDGFRα inhibitor HY-008 is effective against glioblastoma

**DOI:** 10.1101/2022.10.16.512412

**Authors:** Chenxin Xu, Haizhong Feng, Weilin Sun

## Abstract

**Background:** Glioblastoma multiforme (GBM) is the most common primary intracranial malignant tumor in adults, with poor prognosis and high recurrence. Routine treatments of GBM show unsatisfactory efficiency in improving patients’ survival because of limited area of surgical resection and drug resistance. New therapeutic agents are needed to improve GBM treatment efficiency, but the blood-brain barrier (BBB) permeability is a major hurdle. Here, we report HY-008 as a promising therapeutic drug targeting PDGFRα signaling with high BBB permeability and efficient inhibiting effects both in vitro and in vivo.

**Methods:** Through structural modification and medicinal chemistry efforts, HY-007 and HY-008 were developed. The brain and plasma pharmacokinetic profiles of these two compounds were assessed. The inhibitory efficiency of HY-007 and HY-008 on GBM cell survival and PDGFRα signaling were evaluated. The efficacy of HY-007 and HY-008 as a single agent or HY-008 in combination with temozolomide (TMZ) was investigated using transformed mouse astrocyte and glioma stem-like cell (GSC) orthotopic xenograft models.

**Results:** HY-007 and HY-008 both had good brain permeability and desirable PK profiles with mild hERG inhibition, while HY-008 is more brain permeable than HY-007. In vitro, HY-007 and HY-008 both significantly inhibited viability of the established GBM cells with PDGF-A overexpression and transformed mouse astrocytes with PDGF-A/PDGFRα overexpression by targeting the PDGFRα signaling activated Erk1/2 and Akt. In vivo, HY-007 and HY-008 both effectively inhibited orthotopic GBM tumor xenograft growth and prolonged the survival of mice, and HY-008 showed less toxicity and better therapeutic effect. In addition, HY-008 increased sensitivity of TMZ, exhibited treatment efficiency both as a single agent and in combination with TMZ, providing significant survival benefits for GBM tumor xenograft-bearing mice.

**Conclusions:** Our data demonstrate that HY-008 is a promising therapeutic agent in GBM treatment and a combination HY-008 with TMZ could serves as a potential efficient therapeutic option for improving GBM clinical treatment.

## Background

Glioblastoma multiforme (GBM) is the most frequently occurring primary malignant and deadliest form of brain tumor [1]. GBM has a very poor prognosis with a median survival time of less than 15 months, only 27% of patients survive for 2 years, and less than 5% of patients survive for 5 years following diagnosis [2]. Standard therapy for GBM is maximal surgical resection of tumor tissue followed by radiation plus cytotoxic concomitant and adjuvant Temozolomide (TMZ) with at least six TMZ maintenance cycles [3]. After the treatment, GBM always recurs and turns out to be fatal rapidly since the current therapies are generally not very efficient and could hardly increase patients’ survival time [4]. In the past four decades, only four drugs were approved by the FDA for GBM clinical treatment while three of them including TMZ are alkylating agents [5], which lack pertinence of oncogene and individual disease specificity. Because of all the above, GBM remains to be one of the most difficult-to-treat cancer types and there is an immense unmet medical need to develop more effective target therapies for this fatal disease.

In GBM, numerous receptor tyrosine kinases (RTKs) play roles in cancer cell growth, survival, migration, invasion, and angiogenesis [6]. Among these RTKs, amplification of platelet-derived growth factor receptor (PDGFR) is the second most frequent alteration after *EGFR* [7]. *PDGF* family expression and genetic profiles can be prognostic factors and valid therapeutic targets for GBM [8,9]. The proneural subtype of GBM is least responsive to the standard treatment, with a high frequency of mutations in *TP53* in conjunction with overexpression of PDGFRα [7]. Moreover, genetic alterations of components in the PDGFRα-PI3K-AKT signaling pathway are found in over 70% of GBM, and overexpression of PDGFRα, together with the loss of ARF induces GBM through PDGFRα-PI3K-AKT pathway [10,11]. Tumor microenvironment, such as hypoxia is one of the major extrinsic factors contributing to GBM resistance [12]. Hypoxic conditions have been shown to decrease the response of GBM cells to the standard TMZ treatment [13]. Recently, it was reported that hypoxia-induced overexpression of PDGFRα, could further promote tumor invasion and angiogenesis through Akt activation in GBM [14]. Additionally, depletion of PDGFRα diminished glioblastoma stem cell features through PDGFRα/Stat3/Rb1 regulatory axis [15]. Taken together, targeting PDGFR signaling pathway could serve as an effective therapeutic avenue for the treatment of GBM.

Blood-brain barrier (BBB) permeability is a major hurdle for GBM drug development and inadequately delivered concentration of drugs to the brain is one of the main reasons for the failures of most target therapies [16]. Almost all the PDGFR inhibitors attempted for GBM are repurposed drugs that are originally indicated for peripheral cancers, such as Dasatinib, Sunitinib, and Imatinib. Being a substrate of efflux transporter for these compounds may contribute to their poor brain penetration [17-19]. Importantly, contrary to the popularized view that BBB is uniformly disrupted in all GBM patients, overwhelming clinical evidence supports a clinically significant tumor burden with an intact BBB in all GBM [20]. Inferior and inconsistent brain drug concentration presents a great challenge for GBM therapeutics development. Therefore, the development of BBB permeable PDFGR inhibitors is critical for success in GBM treatment.

Here we report a novel PDGFR inhibitor as a promising therapeutic drug for GBM that can deliver and retain pharmacological concentration of the compound in the brain with desirable PK/toxicity profiles as well as shown in vitro/in vivo efficacy against PDGFRα-activated GBM.

## Materials and Methods

### Cell lines

Mouse *Ink4a*/*Arf* ^-/-^ astrocytes and human LN444 cells with overexpression of exogenous PDGF-A and/or PDGFRα were established and characterized as we previously described [21]. All glioma and GSC cells were maintained and cultured as we previously described [21]. Cell lines in this study were recently authenticated using STR DNA fingerprinting (Shanghai, China) and all cultures were routinely tested for mycoplasma contamination.

### Antibodies and reagents

The following antibodies were used in this study: anti-PDGFRα (C-20), anti-phospho-PDGFRα (Y754), anti-phospho-Erk (E-4), and anti–β-actin (I-19) antibodies (Santa Cruz Biotechnology, Dallas, TX, USA); anti-p44/42 MAP Kinase (#9102), anti-phospho-Akt (S473, #4060), and anti-Akt (#9272) antibodies (Cell Signaling Technology, Danvers, MA, USA); anti-CD31 antibody (MEC13.3, BD PHARMIGEN). The secondary antibodies were from Jackson ImmunoResearch Laboratories (West Grove, PA, USA). Peroxidase blocking reagent was from DAKO (Carpinteria, CA, USA); AquaBlock was from East Coast Biologics, Inc (North Berwick, ME, USA). Cell culture media and other reagents were from Invitrogen (Carlsbad, CA, USA), Sigma-Aldrich, or Peprotech (Rocky Hill, NJ, USA).

### Kinase assays

Generally, kinase-tagged T7 phage strains were prepared in an *E. coli* host derived from the BL21 strain (KINOMEscan™ by Eurofins Discovery). E. coli were grown to log-phase and infected with T7phase and incubated with shaking at 32°C until lysis. The lysates were centrifuged and filtered to remove cell debris. The remaining kinases were produced in HEK-293 cells and subsequently tagged with DNA for qPCR detection. Streptavidin-coated magnetic beads were treated with biotinylated small molecule ligands for 30 min at room temperature to generate affinity resins for kinase assays. The liganded beads were blocked with excess biotin and washed with a blocking buffer to remove unbound ligands and reduce non-specific binding. Binding reactions were assembled by combining kinases, liganded affinity beads, and test compounds in a 1x binding buffer. Test compounds solution was sequentially diluted with a duplicate for each concentration point. After incubation, wash, and detection, the binding constants were calculated with a standard dose-response curve using the Hill equation. Curves were fitted using a non-linear least square fit with the Levenberg-Marquardt algorithm.

### Pharmacokinetic studies

Specific-pathogen free Sprague Dawley male rats (Medicilon colony: 999M-017; source: Sino-British SIPPR/BK Lab Animal Ltd, Shanghai) were administered by oral gavage with a single dose of tested compounds (50 mg/kg). The animals were fasted overnight (10-14 hrs) and the food supply to the animals was resumed 4 hrs post-dose. The brain tissues and plasma samples were collected at 8 indicated time points post-dose (0.15 h, 0.5 h, 1 h, 2 h, 4 h, 6 h, 8 h, and 24 h). Three animals were treated for each time point. After processing, the samples were analyzed by UPLC-MS/MS (TQ5500, triple quadrupole). The analytical results were confirmed using quality control samples for intra-assay variation. The accuracy of > 66.7% of the quality control samples is between 80-120% of the known values. A standard set of parameters were calculated using noncompartmental analysis modules in FDA certified pharmacokinetic program Phoenix WinNonlin 7.0 (Pharsight, USA). The ratio of brain tissue to plasma was calculated by the concentration in brain tissue/the concentration in plasma.

### Patch-clamp hERG Cellular Assay

Following the current International Conference on Harmonization (ICH) Harmonized Tripartite Guideline and generally accepted procedures, the compounds were tested on HEK 293 cell line stably expressing hERG (human ether-à-go-go-related gene) potassium channels using a manual patch-clamp system (EPC 10 USB PatchMaster software, HEKA Elektronik). The effects on the electric current passing through hERG were measured in duplicate and the concentration-response relationship for hERG current inhibition by these compounds was defined using the manual patch-clamp technique. Electronic raw data was generated from manual acquisition software and exported to data analysis software Prism Graphpad. The final IC_50_ was obtained by curve fitting with the data analysis software.

### Western blotting assay

The western blotting assay was performed as we previously described [22]. Briefly, cells were lysed in a buffer (20 mM Tris-HCl, pH 7.5, 150 mM NaCl, 1 mM EDTA, 2 mM Na3VO4, 5 mM NaF, 1%Triton X-100, and protease inhibitor cocktail) at 4°C for 20 min, and then the lysates were centrifuged for 20 min at 12,000 x g to remove debris. Protein concentrations were determined with a BCA protein assay kit (Beyotime Biotechnology, China). Equal amounts of cell lysates were resolved in a 2X SDS lysis buffer and analyzed.

### Cell viability assays

Cell viability assay was performed using a WST-1 assay kit (Roche). Briefly, cells were seeded in triplicate wells of a 96-well microplate (4000 cells/Well for LN444 cell and 2000 cells/Well for *Ink4a*/*Arf* ^-/-^ astrocyte). After 24 h, cells were treated with vehicle (ddH_2_O) or HY-007 (HY-008) from 0.1 to 100 μM for 48 h. Then cell viability was analyzed, and half-maximal inhibitory concentration (IC_50_) was calculated from fitted concentration-response curves obtained from at least three independent experiments with GraphPad Prism 9.0 nonlinear regression curve fit.

### Colony formation analysis

For colony formation assay, various cells were dissociated and seeded in 6-well plates (500 cells/well) and maintained in a DMEM medium containing 10% FBS for 2 weeks. The medium was replaced with vehicle (ddH2O) or gradient concentration of HY-007 (HY-008) every 3 days. When a single colony in the control well contained over 50 cells, the plate was removed from the incubator, fixed with 4% paraformaldehyde, and visible colonies were counted after crystal violet staining.

### Tumorigenesis studies

Athymic (Ncr nu/nu) female mice aged 6-8 weeks (SLAC, Shanghai) were used for all animal experiments. All experiments using animals were performed in accordance with a protocol approved by Shanghai Jiao Tong University Institutional Animal Care and Use Committee (IACUC). Mice were randomly divided into 5-6 per group. In total, 1 × 10^5^ *Ink4a/Arf*^-/-^ mAsts or GSC 527 cells were stereotactically implanted into the brain of individual mice. Animals were euthanized when neuropathological symptoms developed. Treatment schemes were described in the corresponding figure legends. Tumor volumes were measured in vivo using luciferase activity after injection of D-luciferin. Mice group allocation, surgery, and assessing the outcome of mice were performed independently by different investigators.

### Immunohistochemistry (IHC) of mouse glioma specimens

Immunohistochemical staining (IHC) was performed as we previously described [23]. Briefly, 6 μM thick OCT-embedded mouse fresh frozen brain sections containing glioma xenograft tumors were separately stained with antibodies against p-AKT (1:100), Akt (1:200), or CD31 (1:1000). After incubated with secondary antibody, images were captured with an Olympus BX53 microscope equipped with an Olympus DP73 digital camera. The percentage of positive cells from five random images per section of mouse brains was calculated by using the formula [A (5 low power fields of positive staining)/ B (total count per 5 fields) × 100]. Statistical analyses were performed using GraphPad Software.

### Statistical analysis

Statistical analyses were performed in a GraphPad Prism version 6.0 for Windows (GraphPad Software Inc., San Diego, CA, USA). Survival analysis was carried out using log-rank tests, and a Mantel-Haneszel approach was used to determine the hazard ratio. A two-tailed Fisher’s exact test was performed to determine if the frequency distribution of the variables were statistically significant. Comparison of treatments was analyzed using One-way ANOVA with Newman-Keuls post-test or a paired two-way Student’s *t*-test as we previously described. *P* values less than 0.05 were considered significant.

## Results

### HY-007 and HY-008 are potent PDGFRα inhibitors

Through structural modification and medicinal chemistry efforts, Sun at Hongyi LLC has developed a series of novel compounds with significantly improved brain permeability, favorable

PK profiles, and a lack of cardiovascular toxicity [24]. After preliminary screening, compounds HY-007 and HY-008 were selected for further investigation as lead PDGFRα inhibitors for the treatment of GBM. The binding affinity to PDGFRα of the synthesized compounds was tested in duplicate through KINOMEscan™ by Eurofins Discovery. The Kd values of HY-007 and HY-008 were 79 and 73 nM, respectively in the binding assay (Figure 1A and 1B).

**Figure 1.**
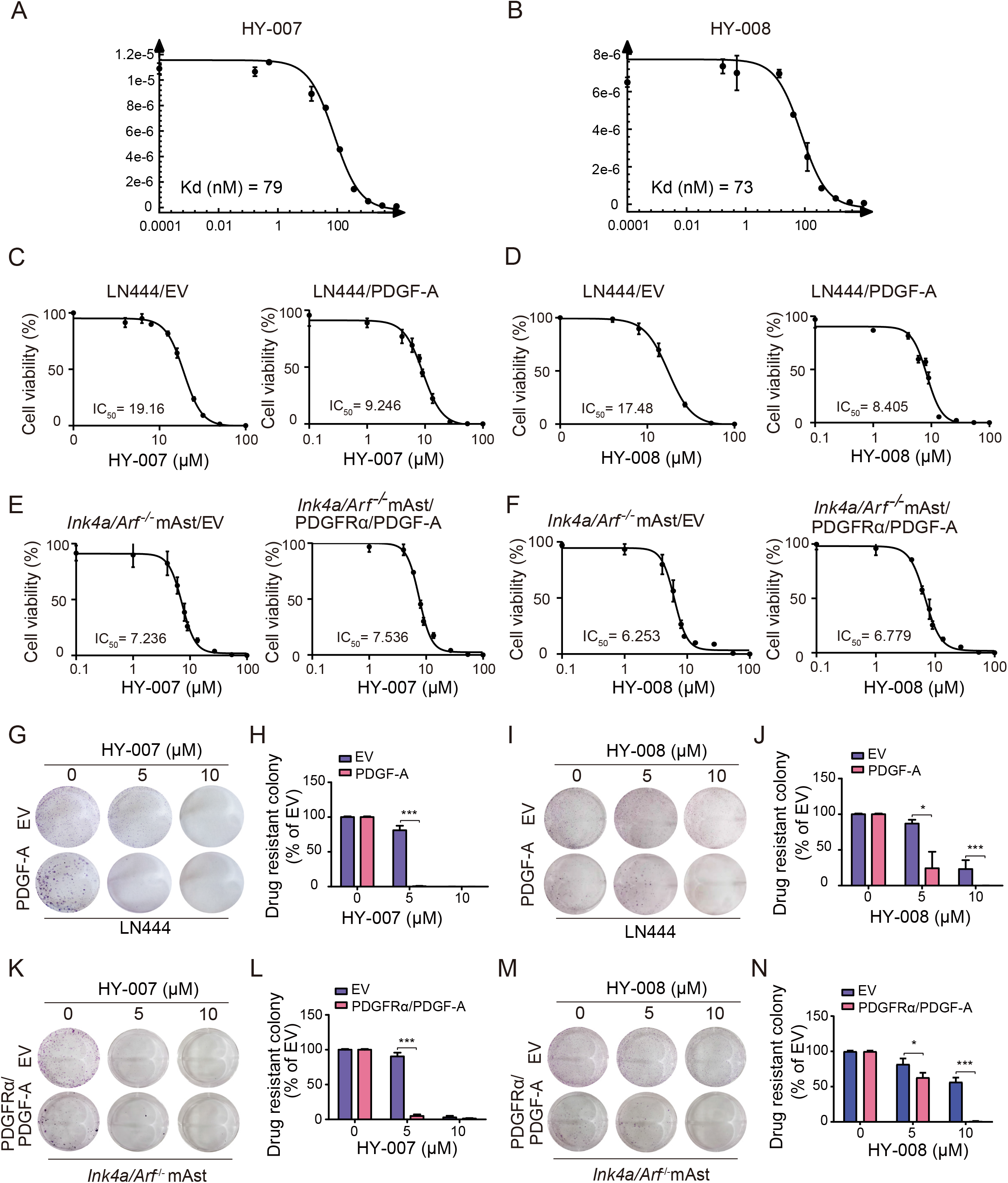
HY-007 and HY-008 are potent PDGFRα inhibitors. **A** and **B**) Binding affinity curve of HY-007 (A) and HY-008 (B) to PDGFRα. The binding affinity to PDGFRα of the synthesized compounds was tested in duplicate through KINOMEscan™ by Eurofins Discovery. **C** and **D**) Viability of LN444 cells with ectopic expression of an empty vector (EV) or PDGF-A at 48 h after treatment with HY-007 (**C**) or HY-008 (**D**). **E** and **F**) Viability of mouse *Ink4a*/*Arf* ^-/-^ astrocytes (*Ink4a*/*Arf* ^-/-^ mAsts) with ectopic expression of an EV or PDGFRα plus PDGF-A at 48 h after treatment with HY-007 (**E**) or HY-008 (**F**). **G** and **H**) Representative images of drug-resistant colony formation in LN444 cells (**G**) or *Ink4a*/*Arf* ^-/-^ mAsts (**H**) at Day 14 post HY-007 treatment. **I** and **J**) Representative images of drug-resistant colony formation in LN444 cells (**I**) or *Ink4a*/*Arf* ^-/-^ mAsts (**J**) on Day 14 post HY-008 treatment. **K** and **L**) Quantification of drug-resistant colony formation in (**I**) and (**J**), respectively. **M** and **N**) Quantification of drug-resistant colony formation in (**K**) and (**L**), respectively. EV, empty vector. *P* value was calculated by one-way ANOVA or two-tailed Student’s *t*-test.

Then, HY-007 and HY-008 were evaluated in cellular assays using human LN444 GBM cancer cells and mouse *Ink4a/Arf*^*-/-*^ astrocytes (*Ink4a/Arf*^*-/-*^ mAsts). *Ink4a/Arf*^*-/-*^ mAsts with both *INK4A* and *ARF* gene deletion are prone to carcinogenesis and generate GBM cancer cells [25]. Both cell lines are responsive to PDGF-A stimulation and up-regulate the expression of downstream molecules such as phosphorylated PDGFRα, ERK1/2, and Akt kinases [21]. As shown in Figures 1C and 1D, compared with the empty vector (EV) control, LN444 GBM cells overexpressed PDGF-A with features of PDGFRα activation and tumorigenicity are more responsive to the treatment of HY-007 and HY-008. The IC_50_ of the LN444 GBM cell viability assay for the treatment group was changed about two-fold compared to the control group, suggesting that compounds HY-007 and HY-008 are likely engaged in the target PDGFRα. Interestingly, there is almost no difference in IC_50_ of *Ink4a/Arf*^*-/-*^ mAsts viability assay between the treatment and control groups as shown in Figures 1E and 1F. Similarly, HY-007 and HY-008 significantly inhibited colony formation of LN444 cells with PDGF-A overexpression compared with the control groups (Figure 1G-J). But for *Ink4a/Arf*^*-/-*^ mAsts with PDGFRα/PDGF-A overexpression, HY-007 and HY-008 also showed significant inhibitory effects on cell viability, contrary to the sight difference in IC_50_ (Figure 1K-N). These results indicate that human glioma cells with PDGFR activation are more responsive to HY-007 and HY-008 through their inhibition of PDGFRα.

### HY-007 and HY-008 are brain penetrant

Since blood-brain barrier permeability is critical for GBM drug development, we next carried out in vivo PK studies of these two selected compounds. HY-007 and HY-008 at a single dose of 50 mg/kg were administered orally to Sprague Dawley male rats. Blood and brain tissues were collected in triplicate at 8 time-points after the dose: 0.25 h, 0.5 h, 1 h, 2 h, 4 h, 6 h, 8 h, and 24 h. Drug concentrations in blood and the brain were analyzed by LC-MS/MS. The analytical results of plasma and brain concentrations were confirmed using quality control samples for intra-assay variation (Figures 2A and 2B). The accuracy of >66.7% of the quality control samples is between 80-120% of the known values. PK profile results of HY-007 and HY-008 are listed in Figure 2C. A standard set of parameters including area under the curve (AUC_(0-t)_ and AUC_(0-_∞)), elimination half-life (T_1/2_), maximum plasma concentration (C_max_), and time to reach maximum plasma concentration (T_max_) was calculated using noncompartmental analysis modules in FDA certified pharmacokinetic program Phoenix WinNonlin 7.0 (Pharsight, USA). The ratio of brain to plasma concentration were calculated by the concentration in brain tissue/the concentration in plasma (Supplementary Table 1). Nilotinib is an inhibitor of multiple kinases including PDGFR and it was also evaluated on brain penetration for comparison. Our data show that HY-007 and HY-008 are brain penetrant.

**Figure 2.**
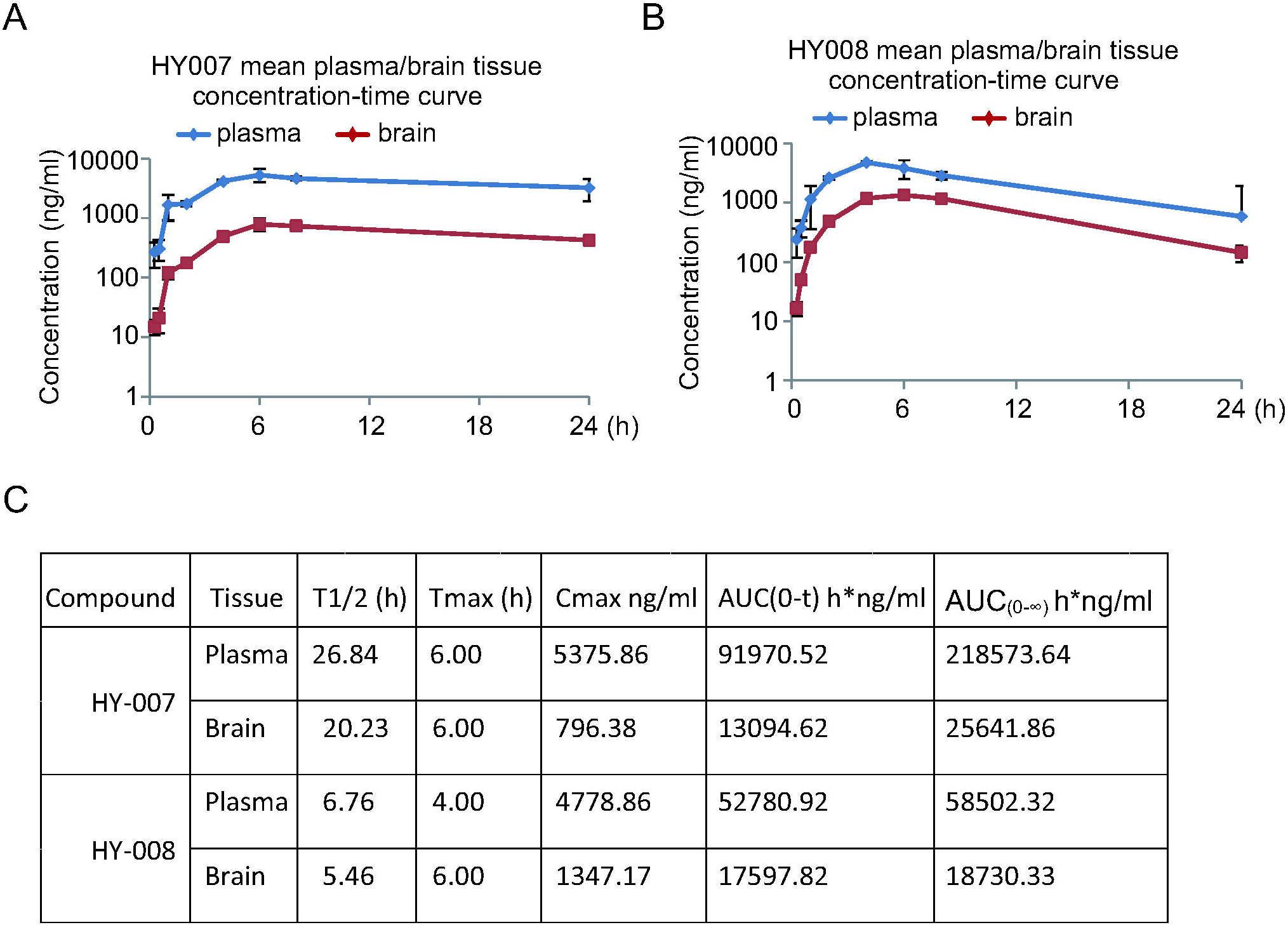
HY-007 and HY-008 are brain penetrant. HY-007 (**A**) and HY-008 (**B**) at a single dose of 50 mg/kg were administered orally to Sprague Dawley male rats. Blood and brain tissues were collected in triplicate at 8 time points after the dose: 0.25 h, 0.5 h, 1 h, 2 h, 4 h, 6 h, 8 h, and 24 h. Drug concentrations in blood and the brain were analyzed by LC-MS/MS. Plasma concentration is in ng/mL and brain tissue concentration is in ng/g.

### HY-007 and HY-008 show weak hERG inhibition

Blockade of the hERG potassium channel can induce cardiac arrhythmia and is one of the main concerns in drug development. Nilotinib has shown mild efficacy in GBM in a phase-2 clinical trial [26]. But as a strong hERG inhibitor with IC_50_ around 0.13 μM, Nilotinib has a black box warning for its QT prolongation toxicity effect [27]. To minimize the potential cardiovascular toxicity liability, HY-007 and HY-008 were screened in an ICH-compatible hERG potassium channel inhibition assay. HY-007 and HY-008 showed mild inhibition of the hERG channel with IC_50_ of 2.52 and 3.33 μM, respectively(Figures 3A and 3B)These findings support that HY-007 and HY-008 show weak hERG inhibition and potentially low cardiovascular toxicity.

**Figure 3.**
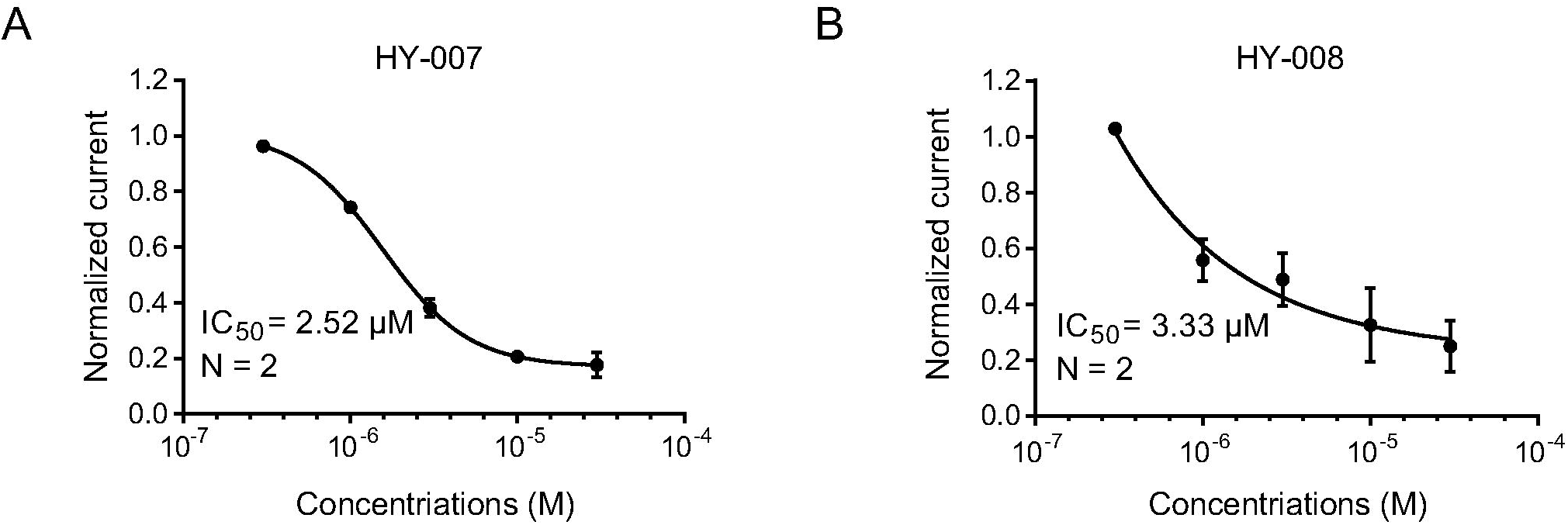
HY-007 and HY-008 show weak hERG inhibition. **A** and **B**) Blocking activity curves on whole cell hERG currents by the patch-clamp hERG whole cell assay. HY-007 (**A**) and HY-008 (**B**) were investigated at concentrations of 0.3, 1, 3, 10, and 30 μM. IC_50_ values which were fitted by the Hill equation are 2.52 μM and 3.33 μM, respectively. Terfenadine was used as a positive control for IC_50_ determination.

### HY-007 and HY-008 inhibit the activation of PDGFR, Akt, and ERK1/2

*Ink4a/Arf*^*-/-*^ mAsts were treated with HY-007 and HY-008. Compared to EV control, overexpression of PDGFRα/PDGF-A stimulated the phosphorylation of PDGFR, Akt, and ERK1/2 kinases, consistent with previously reported results [21]. In the western blot analyses as shown in Figures 4A and 4B, the activation of phosphorylation of PDGFR, Akt, and ERK1/2 in the PDGFRα/PDGF-A overexpressed *Ink4a/Arf*^*-/-*^ mAsts were further inhibited by HY-007 and HY-008 treatments compared to the vehicle-treated cells. Similar inhibitory effects were also shown in the LN444 cells, to which we applied gradient concentrations of HY-007 and HY-008 (Figures 4C and 4D). These results demonstrate that HY-007 and HY-008 could efficiently inhibit GBM cell growth by targeting PDGFR, Akt, and ERK pathways.

**Figure 4.**
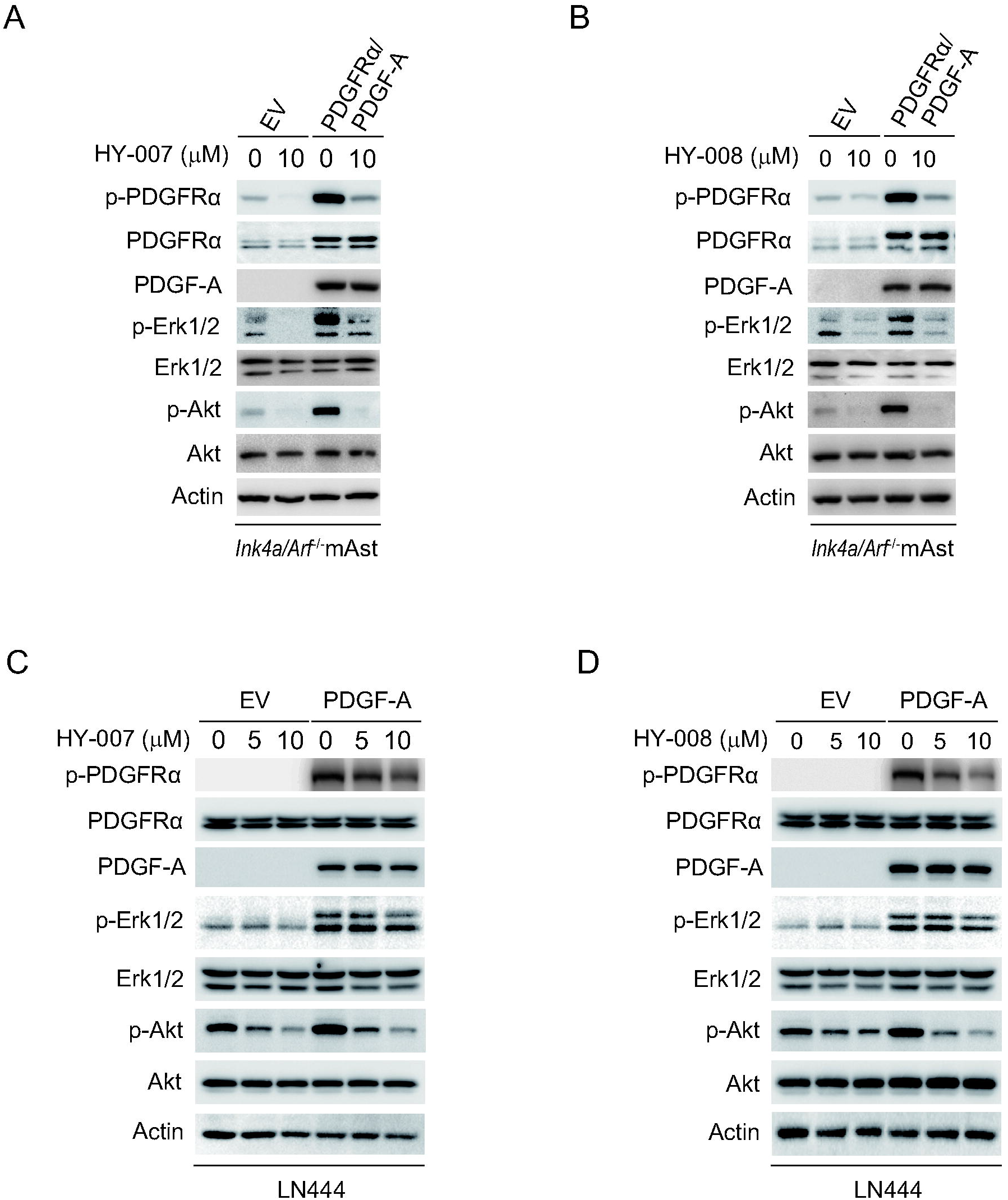
HY-007 and HY-008 specifically inhibit PDGFRα activated Erk1/2 and Akt activity. **A** and **B**) WB assay of effects of HY-007 (**A**) or HY-008 (**B**) on PDGF-A-activated p-PDGFRα, Erk1/2 phosphorylation (p-Erk1/2), and Akt phosphorylation (p-Akt) in *Ink4a*/*Arf* ^-/-^ mAsts overexpressing an empty vector (EV) and PDGFRα plus PDGF-A. **C** and **D**) WB assay of effects of HY-007 (**C**) or HY-008 (**D**) on PDGF-A-activated p-PDGFRα, Erk1/2 phosphorylation (p-Erk1/2), and Akt phosphorylation (p-Akt) in LN444 cells overexpressing an empty vector (EV) and PDGF-A. EV, empty vector.

### HY-008 inhibits PDGFRα-driven glioma tumor growth as a single agent

Since HY-007 and HY-008 showed similar efficacy in cellular assays while HY-008 is more brain permeable than HY-007, we performed in vivo studies with both HY-007 and HY-008 but selected HY-008 for further studies and analysis. An orthotopic xenograft model in immunodeficient mice was used to evaluate in vivo antitumor efficacy of both drugs. In a preliminary animal study, *Ink4a/Arf*^*-/-*^ mAsts with stable expression of PDGFRα/PDGF-A and luciferase were stereotactically transplanted into the brains of the mice. HY-007 and HY-008 were administered by I.P. after randomization by bioluminescence imaging (BLI) flux values to ensure that tumors were of equal size in the treatment and control groups. On day 4, vehicle, HY-007 or HY-008 at 30 mg/kg and 60 mg/kg were given for 5 consecutive days, after two days off, followed by 5 more consecutive days (Figure 5A). The tumor sizes in HY-007 treatment groups were significantly decreased compared to control cohorts evaluated by bioluminescence imaging (BLI), as shown in Figures 5B and 5C, and the survival rates of 30 mg/kg of HY-007 cohorts were significantly improved compared with the control cohort (*P* < 0.01), as shown in Figure 5D. However, the survival rates of 60 mg/kg of HY-007 cohorts showed little difference from control cohorts, possibly because of the high toxicity of the high dose of HY-007 in vivo (Figure 5D). The median survival days for these three groups were 19 days for control, 27 days for HY-007 at 30 mg/kg, and 18 days for HY-007 at 60 mg/kg. In HY-008 treatment groups, the tumor sizes also showed significantly decreased compared to control groups. While the difference between 30 and 60 mg/kg of HY-008 treatment cohorts was not significant due to inadequate sample size (n = 5), HY-008 still showed inhibition of glioma tumor growth in a dose-dependent manner (Figures 5E and 5F). Different from HY-007, the survival rates of 30 and 60 mg/kg of HY-008 cohorts were both significantly improved compared with the control cohort (*P* < 0.01) (Figure 5G), suggesting that HY-008 had more potential of being applied in clinical GBM treatment. The median survival days for these three groups were 19 days for control, 27 days for HY-008 at 30 mg/kg, and 31 days for HY-008 at 60 mg/kg. Besides, IHC analyses confirmed the anti-tumor efficacy of HY-008 evidenced by decreased p-Akt and tumor vascularity marker CD31 dose-dependently as shown in Figures 5H and 5I. The results above indicate that HY008 may serve as an efficient therapeutic agent for GBM patients with PDGFRα amplification.

**Figure 5.**
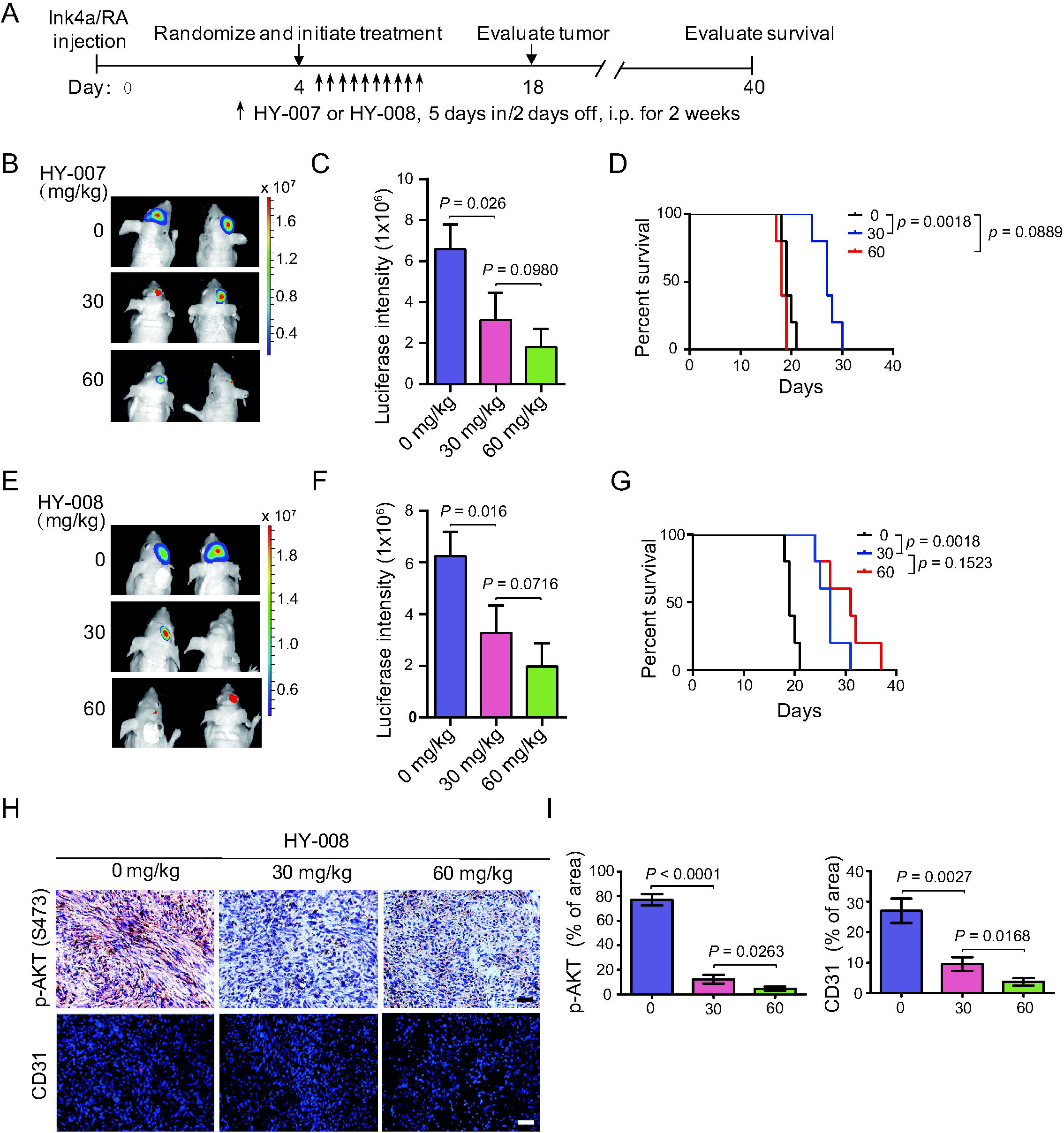
HY-007 and HY-008 can serve as a single agent to effectively inhibit PDGFRα-driven glioma tumor growth. **A**) Treatment scheme for the evaluation of in vivo efficacy of HY-007 and HY-008 in *Ink4a*/*Arf* ^-/-^ mAsts xenografts. Animals were treated with HY-007 or HY-008 of the indicated doses from Monday to Friday within two weeks. **B** and **C**) Representative images (**B**) at Day 18 and quantitation of BLI (**C**) of *Ink4a*/*Arf* ^-/-^ mAsts xenografts with ectopic expression of PDGFRα plus PDGF-A from HY-007 treated and control mice. **D**) Kaplan-Meier survival analysis of animals with *Ink4a*/*Arf* ^-/-^ mAsts tumors (n=5 or 6 per group). **E** and **F**) Representative images (**E**) at Day 18 and quantitation of BLI (**F**) of *Ink4a*/*Arf* ^-/-^ mAsts xenografts with ectopic expression of PDGFRα plus PDGF-A from HY-008 treated and control mice. **G)** Kaplan-Meier survival analysis of animals with *Ink4a*/*Arf* ^-/-^ mAsts tumors (n=5 or 6 per group). **H)** Representative images of IHC analysis of xenograft in (**E**) using anti-p-Akt and anti-CD31 antibodies. **I)** Quantitation of p-Akt and CD31 positive cells. In **H**, scale bars, 25 μm. *P* value was calculated by two-tailed Student’s *t*-test or log-rank analysis.

### HY-008 alone and in combination with TMZ extends the survival of GBM-bearing mice

Next, we evaluated the anti-tumor efficacy of HY-008 in the expanded animal studies using two different glioma cell lines. TMZ, an alkylating agent, is used in the standard treatment of GBM. To investigate whether HY-008 could enhance the chemosensitivity of TMZ, xenograft model mice bearing *Ink4a/Arf*^*-/-*^ mAsts that expressed PDGFRα/PDGF-A or human GSC 157 tumor cells with highly expressed PDGFRα and p-PDGFRα were treated with TMZ (5 mg/kg), HY-008 (60 mg/kg), combination of HY-008 (60 mg/kg) with TMZ (5 mg/kg), and vehicle for comparison. After randomization by BLI, I.P. administration of HY-008 and TMZ was initiated. On day 18 (*Ink4a/Arf*^*-/-*^ mAsts) or day 50 (GSC 157) post-implantation, xenograft tumor sizes were measured by BLI (Figures 6A and 6E). HY-008 alone or HY-008 in combination with TMZ significantly inhibited tumor growth compared to TMZ treatment and vehicle cohorts (Figure 6B, 6C, 6F, and 6G). HY-008 alone or HY-008 in combination with TMZ also significantly extends the survival of GBM xenograft mice (Figure 6D and 6H), suggesting that HY-008 increases the chemosensitivity of TMZ and synergizes with TMZ to extend the survival of GBM model mice.

**Figure 6.**
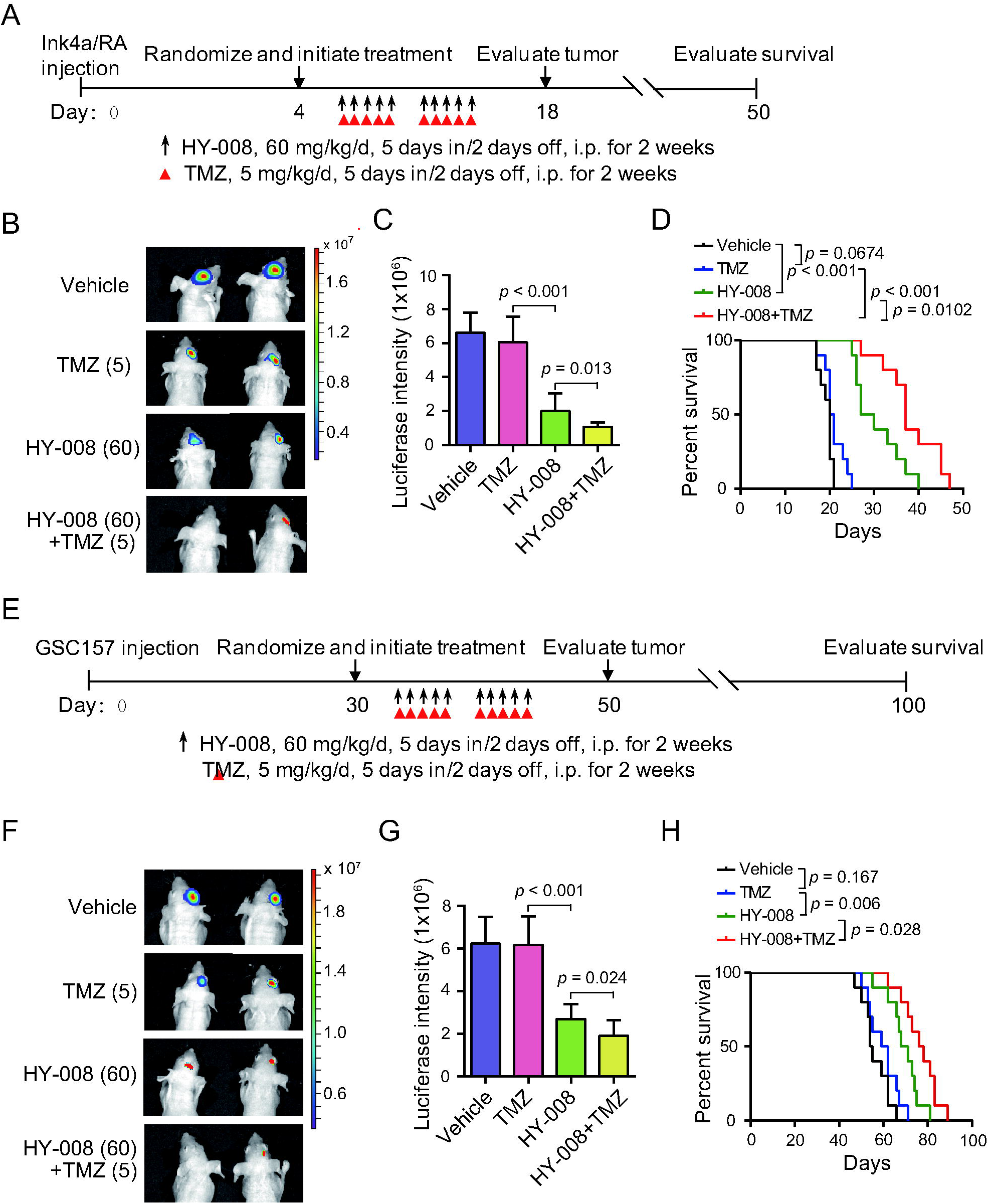
HY-008 in combination with TMZ prolonged the survival of GBM-bearing mice. **A** and **E**) Treatment scheme for the evaluation of in vivo efficacy of HY-008 in combination with TMZ in *Ink4a*/*Arf* ^-/-^ mAsts (**A**) or GSC 157 (**E**) tumor xenografts. The mice were treated with indicated 60 mg/kg HY-008 with or without 5 mg/kg TMZ from Monday to Friday within two weeks. **B** and **C**) Representative images (**B**) at Day 18 and quantitation of BLI at the indicated date (**C**) of *Ink4a*/*Arf* ^-/-^ mAsts xenografts with ectopic expression of PDGFRα plus PDGF-A from HY-008 in combination with TMZ treated and control mice. **D**) Kaplan-Meier survival analysis of animals with *Ink4a*/*Arf* ^-/-^ mAsts tumors (n=10 per group). **F** and **G**) Representative images (**F**) at Day 50 and quantitation of BLI at the indicated date (**G**) of GSC 157 tumor xenografts from HY-008 in combination with TMZ treated and control mice. **H**) Kaplan-Meier survival analysis of animals with *Ink4a*/*Arf* ^-/-^ mAsts tumors (n=10 per group). *P* value was calculated by two-tailed Student’s *t*-test, one-way ANOVA, or log-rank analysis.

## Discussion

Surgical resection of tumor tissue followed by radiation and TMZ has been used as the standard first-line treatment for GBM for decades but resistance and recurrence lead to poor prognosis for GBM patients. Extensive efforts have been put into developing target therapies that could tackle the responsible resistance mechanism in GBM. PDGFRα overexpression plays an essential role in tumor invasion and angiogenesis through Akt activation in GBM [14]. It was also found that PDGFRα is involved in self-renewal, invasion, and differentiation in glioblastoma stem cells [15]. Proneural subtype GBM, featured with overexpression of PDGFRα, is most resistant to the standard treatment.

Several repurposed PDGFR inhibitors were tested for GBM. However, the poor blood-brain barrier permeability of these drugs is blamed for the failure of these clinical trials [18-20]. Nilotinib, the second generation of BCR-ABL kinase inhibitor, also a PDGFR inhibitor, showed limited efficacy in recurrent GBM enriched for PDGFRα in a phase II trial [26]. In a different trial for Parkinson’s disease with a single dose of Nilotinib, blood and CSF samples from patients were collected 2 h after administration. An average of 0.5-1% Nilotinib in the CSF as a free drug related to the concentration in plasma is detected [28]. In our PK studies, the ratio of brain to plasma concentration ranged from 1.85-7.66% for Nilotinib, suggesting the measured ratio value in our PK studies is in line with the actual brain permeability observed in patients. The ratio of brain to plasma concentration for HY-008 ranges from 6.71-40.72%. HY-008 also showed a favorable kinetics profile. The ratio of the concentration of brain to plasma for HY-008 keeps increasing until 8 h, suggesting that HY-008 continuously penetrates the brain and has longer retention in the brain. These data demonstrated that HY-008 had good brain permeability and a desirable PK profile. Additionally, different from the potent hERG blocking activity and QT prolongation toxicity for Nilotinib, HY-008 showed mild hERG inhibition, indicating a lower toxicity profile compared to Nilotinib.

In cellular assays, human GBM cells overexpressed PDGF-A with features of PDGFRα activation are more responsive to HY-008. HY-008 inhibits downstream phosphorylation of ERK1/2 and Akt promoted by PDGFRα overexpression. Inhibition of the PDGFRα-ERK1/2 signaling pathway reduces the proliferation and migration of glioma cells [29]. Additionally, suppression of PDGFRα decreases tumor invasion and angiogenesis through inhibition of Akt signaling activation [14]. These anti-tumor results were confirmed by IHC data in the *Ink4a/Arf*^*-/-*^ mAsts orthotopic xenograft animal studies in which HY-008 treatment decreased p-Akt and tumor angiogenesis marker CD31 in a dose-dependent manner. In two xenograft model mice bearing *Ink4a/Arf*^*-/-*^ mAsts and human GSC 157 tumor cells, HY-008 alone and in combination with TMZ significantly decreases tumor size and extended the survival of GBM-bearing survival mice.

Despite decades of research into the biology and treatment of GBM, to date the outcome for target molecular therapies of GBM remains poor due to several challenges: 1) insufficient brain penetration; 2) high degree of inter- and intra-tumoral heterogeneity in GBM; 3) plasticity in GBM tumor cells [30]. Moreover, enhanced toxicity from the combination of multiple target therapies requires suboptimal dosing for each drug. To overcome these challenges, it is more promising to develop a single brain-permeable precision therapy that could target multiple relevant oncogenic-driven signaling pathways at the same time in a multiple-headed missile mode. A combination of a multiple-targeting kinase inhibitor with a standard cytotoxic alkylation agent could potentiate efficiency of GBM treatment and reduce resistance and recurrence of standard treatment.

Hereby, we reported the identification of a potent brain penetrant PDGFR inhibitor, HY-008, which showed a desirable PK/toxicity profile and promising efficacy against GBM in both in-vitro and in-vivo studies. HY-008 specifically targets PDGFRα and inhibits the activation of Erk1/2 and Akt signaling pathways. Some PDGFR inhibitor, like Sorafenib, fails to enhance the chemosensitivity of TMZ against GBM cell lines [31]. Notably, HY-008 increases the chemosensitivity of TMZ in GBM animal model studies. A combination of HY-008 with TMZ significantly improved anti-tumor efficacy compared to monotherapy with TMZ, which provides a validated rationale for a combination therapeutic approach and warrants further investigations in future clinical studies.

## Conclusion

In summary, we reported that a novel brain penetrant PDGFRα inhibitor, HY-008, as a single agent or in combination with TMZ, inhibited tumor growth and extended survival of two different orthotopic xenograft glioblastoma animal models. These results support that HY-008 is a promising clinical candidate for GBM treatment.

## Declarations

### Consent for publication

All authors have agreed to publish this manuscript.

### Availability of data and materials

The data in the current study are available from the corresponding author upon request.

### Authors’ contributions

HF and WS designed and supervised the project. CX performed cellular assays and animal studies. CX, HF, and WS interpreted and reviewed the data. WS and HF wrote and edited the manuscript.

