## Supplemental Table 1 for "A novel brain penetrant PDGFRα inhibitor HY-008 is effective against glioblastoma"

**Supplementary Table 1** Mean Brain/plasma ratio of Nilotinib, HY-007, and HY-008

| Compound | 0.25 h | 0.5 h | 1 h | 2 h | 4 h | 6 h | 8 h | 24 h |
| --- | --- | --- | --- | --- | --- | --- | --- | --- |
| Nilotinib | 0.0185 | 0.0294 | 0.0322 | 0.0411 | 0.0440 | 0.0766 | 0.0594 | 0.0323 |
| HY-007 | 0.058 | 0.0799 | 0.0777 | 0.1004 | 0.1173 | 0.1485 | 0.1588 | 0.1429 |
| HY-008 | 0.0671 | 0.1374 | 0.1583 | 0.1825 | 0.2465 | 0.3481 | 0.4072 | 0.2595 |
